# Tolerance to Lung Infection in TWIK2 K^+^ Efflux Mediated Macrophage Trained Immunity

**DOI:** 10.1101/2025.05.25.655979

**Authors:** Josh Thompson, Yufan Li, Yuanling Song, Mohammad Anas, Jaewon Cho, Ki-Wook Kim, Asrar B. Malik, Jingsong Xu

## Abstract

Lung macrophages such as alveolar macrophages (AMφ) are essential for innate immune function in the lungs. It is now apparent that macrophages can be trained to become better at attacking infections. Although trained immunity is thought to result from metabolic and epigenetic reprogramming, the underlying mechanisms remain unclear. Here, we report that macrophages can be trained by extracellular ATP, which is ubiquitously released during inflammation. ATP ligates the canonical Purinergic Receptor 2 subtype X7 receptor (P2X7) to mediate endosomal Two-pore domain Weak Inwardly rectifying K^+^ channel 2 (TWIK2) translocation into the plasma membrane (PM). This enables the cells to transit to a ‘ready’ state for microbial killing in two directions: first, K^+^ efflux via PM-TWIK2 induces NLRP3 inflammasome activation, which further activates metabolic pathways; second, upon bacterial phagocytosis, PM-TWIK2 internalizes into phagosome membrane with proper topological orientation, where TWIK2 mediates K^+^ influx into phagosomes to control pH and ionic strength favoring bacterial killing. Therefore, the enhanced association of TWIK2 in phagosomal and plasma membranes signaled by danger-associated molecular patterns (DAMPs), such as ATP, mediates trained immunity in macrophages and enhances microbiocidal activity.

## Introduction

Immune responses are classically divided into innate immune responses, which react rapidly and nonspecifically upon encountering a pathogen, and adaptive immune responses, which are slower to develop and show a measured and specific response that is the result of acquired immunological memory (Netea et al., 2016). Although innate immunity is considered a non-specific, short-lived phenomenon as opposed to adaptive immunity, which is long-lived and highly specific, recent studies have shown that innate immunity can display adaptive immunity characteristics after challenge with pathogens or their products (Netea et al., 2016). Trained immunity mediates protection against heterologous infections and is thought to be due to epigenetic and functional reprogramming of innate immune cells (Netea and Joosten, 2018). However, trained immunity also exhibits maladaptive effects in chronic inflammatory conditions such as cardiovascular diseases and autoinflammatory syndromes (Netea and Joosten, 2018).

Lung resident macrophages such as AMφ are key effector cells of the innate immune response in the lungs, which not only play a key role in killing bacterial pathogens through processes such as phagocytosis or secretion of antimicrobial peptides, but also play an important role in the restoration of homeostasis (Allard et al., 2018). The efficiency of bacterial killing is an essential determinant of the ability to resolve lung inflammation and injury as seen in patients with acute lung injury (ALI) and acute respiratory distress syndrome (ARDS) (Sevransky et al., 2009). Being located at the interface of the airways and the environment, lung AMφ exhibit a distinctive cellular profile that tightly regulates their activation state to avoid excessive inflammation (Bain and MacDonald, 2022). Due to their unique properties, lung AMφ are important candidates for understanding trained immunity. Our current knowledge about innate immune memory responses of lung macrophages and the underlying mechanisms remains limited.

NLRP3 inflammasome is a key determinant of acute immune responses as seen in ALI/ARSD (Grailer et al., 2014; Sefik et al., 2022; Swanson et al., 2019). Activation of NLRP3 inflammasome complex is a multi-step process involving the assembly of key proteins and activation of caspase-1, which cleaves pro-Interleukin-1β (pro-IL-1β) to release the active form of this inflammatory cytokine (He et al., 2016; Swanson et al., 2019). However, little is known about the triggers that initiate the activation of NLRP3 complex. An essential mechanism of NLRP3 assembly is the efflux of K^+^ (Franchi et al., 2007; Munoz-Planillo et al., 2013; Petrilli et al., 2007) through the PM bound K^+^ channel TWIK2 (Di et al., 2018; Enyedi and Czirjak, 2010). The efflux of K^+^ generates regions of low intracellular K^+^, which promote a conformational change of inactive NLRP3 to facilitate NLRP3 assembly and activation (Tapia-Abellan et al., 2021). TWIK2 mediated K^+^ efflux thus serves as a checkpoint for the initiation of trained immunity as well as maladaptive inflammatory signaling mediated by NLRP3 (Di et al., 2018). This function of TWIK2 in initiating macrophage trained immunity remains uncertain.

Our previous work outlined a mechanism wherein extracellular ATP induces Rab-11 mediated outward translocation of the TWIK2 K^+^ efflux channel (Huang et al., 2023). Membrane-associated TWIK2 was sufficient to drive NLRP3-activating potassium efflux, significantly enhancing inflammatory innate immune function (Di et al., 2018; Huang et al., 2023). Here, we show that TWIK2 after PM insertion (induced by ATP) is re-internalized into macrophages upon infection, while remaining associated with the phagosome membrane. TWIK2 is localized in the phagosome membrane and increases phagosome ionic strength by transporting K^+^ from the cytosol into the phagosome, which favors phagosomal protease activation thus enhancing bactericidal activity.

Therefore, TWIK2 functions both at the PM and in phagosomes to optimize bacterial killing and the channel can be modulated to promote macrophage training. The regulatable association of TWIK2 with the PM and phagosomes triggered by ATP mediates trained immunity in macrophages yet avoids excessive inflammation.

## Results

### TWIK2 is required for ATP induced macrophage training

Lung AMφ are exposed to inhaled particles and microbes and thus represent ideal cells to study trained immunity. Since ATP is widely appreciated as a ubiquitous DAMP (Coutinho-Silva and Savio, 2021) that can activate NLRP3 via P2X7-TWIK2 signaling in macrophages (Di et al., 2018; Huang et al., 2023), we investigated the role of ATP in induction of immune memory. WT, *Twik2^−/−^*, *P2x7^−/−^*, and *Nlrp3^−/−^* mice were intranasally exposed to ATP (40μL 1mM, i.n.) and AMφ were isolated on day 7 to determine acquisition of training. The cells were further challenged with GFP-expressing *Pseudomonas aeruginosa* (GFP-*PA*, **Fig. 1A**). To assess the microbial killing activity, we measured the survival of phagocytosed *PA* within AMφ that had been exposed to extracellular ATP and those that had not, as well as the residual bacterial load. The survival of *PA* from ATP-treated AMφ was markedly reduced as compared with controls, indicating augmented microbial killing ability of these ATP-treated AMφ (**Fig. 1B**). Similar results were observed in bone marrow-derived macrophages (BMDM) trained with ATP *in vitro* following full differentiation (**Fig. 1C**). Importantly, bactericidal activity remained unaffected in AMφ or BMDM from TWIK2, P2X7 or NLRP3 null mice (**Fig. 1B**), indicating a crucial role of P2X7-TWIK2-NLRP3 signaling in ATP-mediated AMφ training. These findings support a model that ATP functions through TWIK2 channel to activate training. Indeed, ATP-trained mice had a better survival rate after being infected intratracheally with *PA*, as compared with untrained controls (**Fig. 1D**). The induction of memory was not observed in AMφ treated with other DAMPs such as NAD (0.5 mM, **Fig. 1E**) (Adriouch et al., 2012) highlighting the importance of ATP signaling.

**Figure 1.**
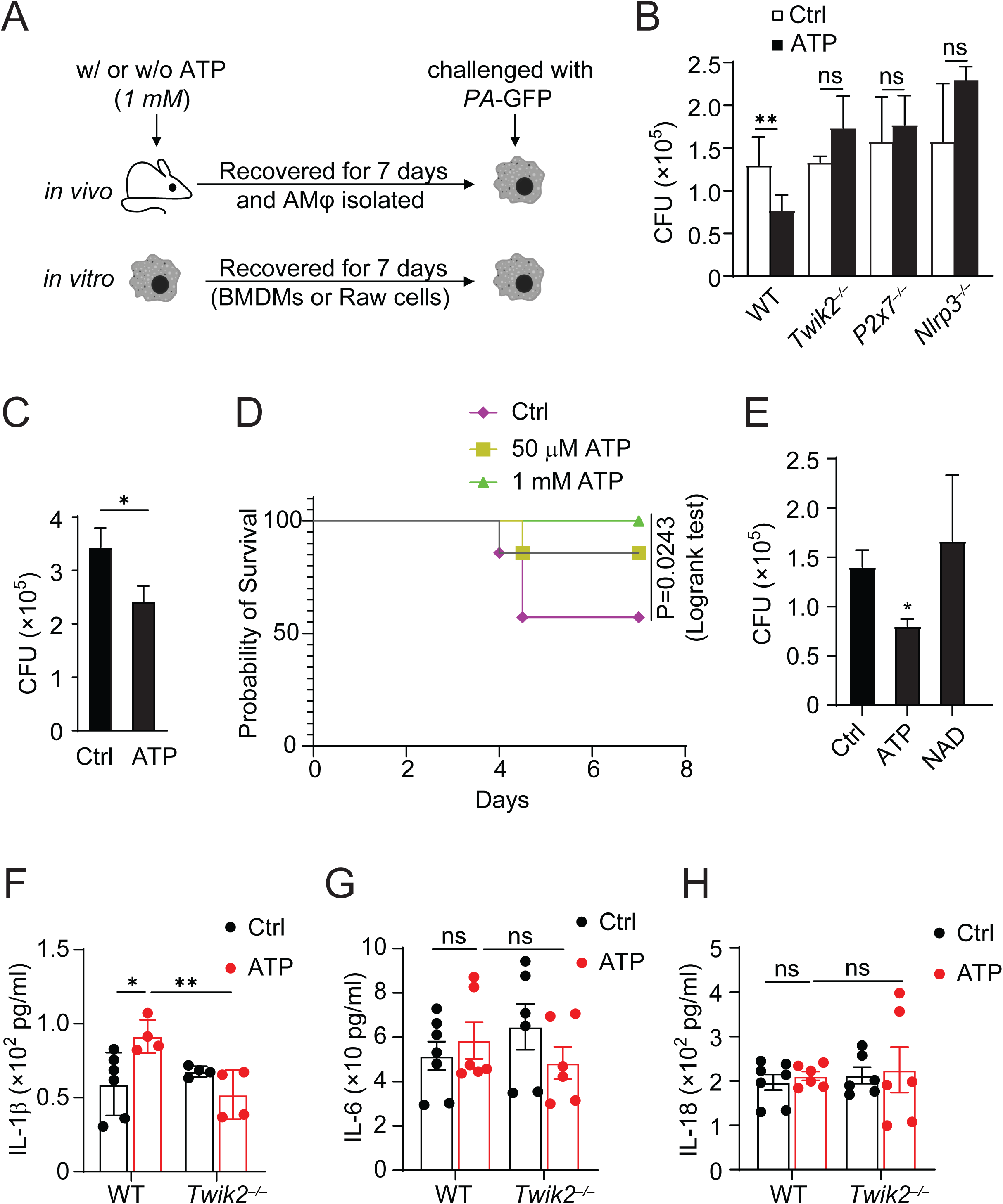
Central role of TWIK2 in triggering ATP-induced macrophage training of bacterial killing. **A.** Diagram of the ATP-induced AMφ training model in vivo. WT or KO mice were exposed to ATP (40 μL at 1mM, i.n.) and AMφ were isolated on day 7 to determine acquisition of training. For in vitro training, BMDMs or RAW cells were trained with or without ATP (1mM) and recovered for 7 days. **B.** AMφ subjected to training as described in A were incubated with live PA-GFP (MOI:1), and assessed for bacterial survival (CFU), n = 3. **C.** BMDMs subjected to in vitro training (1mM ATP for 7 days as described in A) were incubated with live PA-GFP (MOI:1) and were assayed for PA survival (CFU). **D.** Survival of control or ATP pretrained mice after receiving 40mL of 5×10^6^ CFU of *P. aeruginosa* intranasally instilled, n = 7. **E.** Comparison of bacterial load clearing activity of AMφ trained with various DAMPs: ATP, or NAD. ATP showed the most dramatic response. **F-H.** Changes in concentrations of IL1-β (F), IL-6 (G), and IL-18 (H) in ATP-trained AMφ after PA infection. ∗, p<0.05, ∗∗, p<0.01, data are from three independent experiments (mean and s.e.m.). ELISA data (F,G,H) includes measurements from 2-3 technical replicates from each group.

The ATP-trained AMφ also produced greater amount of pro-inflammatory IL-1β as compared to control AMφ after PA exposure, as determined by ELISA (**Fig. 1F**), and this increment was not observed in TWIK2 null AMφ (**Fig. 1F**). Other cytokines such as IL-6 and IL-18 were not significantly changed before and after ATP training (**Fig. 1G, H**). It is worth noting that the populations of immune cells such as neutrophils, monocytes, AMφ and eosinophils were not significantly altered after ATP treatment in lungs (**Fig. 2A-D**), indicating that ATP-induced training may not require newly recruited immune cells into the lungs.

**Figure 2.**
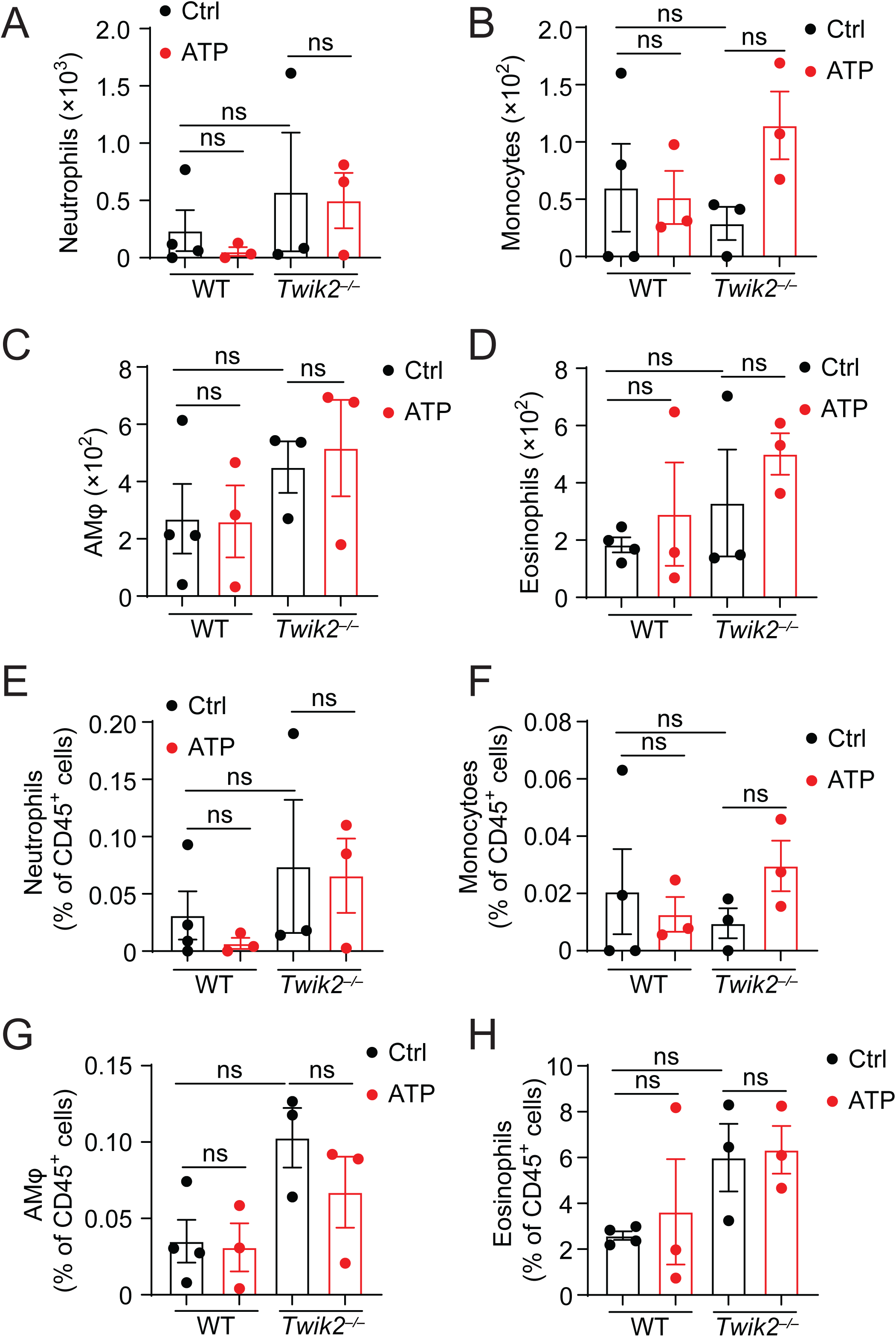
Immune cell counting in the lungs before and after ATP training. **A-D.** Immune cells counting before and after ATP training in lung tissue: neutrophils (A), eosinophils (B), monocytes (C) and AMφ (D). Data are from 3-4 (WT, ctrl) independent experiments (mean and s.e.m.).

### ATP induces endosomal TWIK2 PM translocation in macrophages

We hypothesized that TWIK2 translocation to plasma membrane (PM) and phagosome determines its role in macrophage training, therefore, we studied TWIK2 localization dynamics by imaging TWIK2 PM translocation using TWIK2-GFP. We observed in RAW264.7 cells that TWIK2 translocated to the PM within 2 min post-ATP (**Fig. 3A**); this observation was confirmed by membrane fractioned TWIK2 (**Fig. 3B**). After ATP training, fluorescence recovery after photobleaching (FRAP) demonstrated a significantly reduced mobile fraction for membrane bound TWIK2, indicating that the diffusion of membrane-translocated TWIK2 is indeed constrained by the membrane (**Fig. 3C**). Surprisingly, PM TWIK2 could be positioned at the PM for as long as 6 days following ATP activation (**Fig. 3D**). These data suggest PM association may sustain functionally relevant macrophage training. Interestingly, LPS itself showed no such effect on TWIK2 PM association (**Fig. 3D**) suggesting the importance of ATP signaling in activating the process.

**Figure 3.**
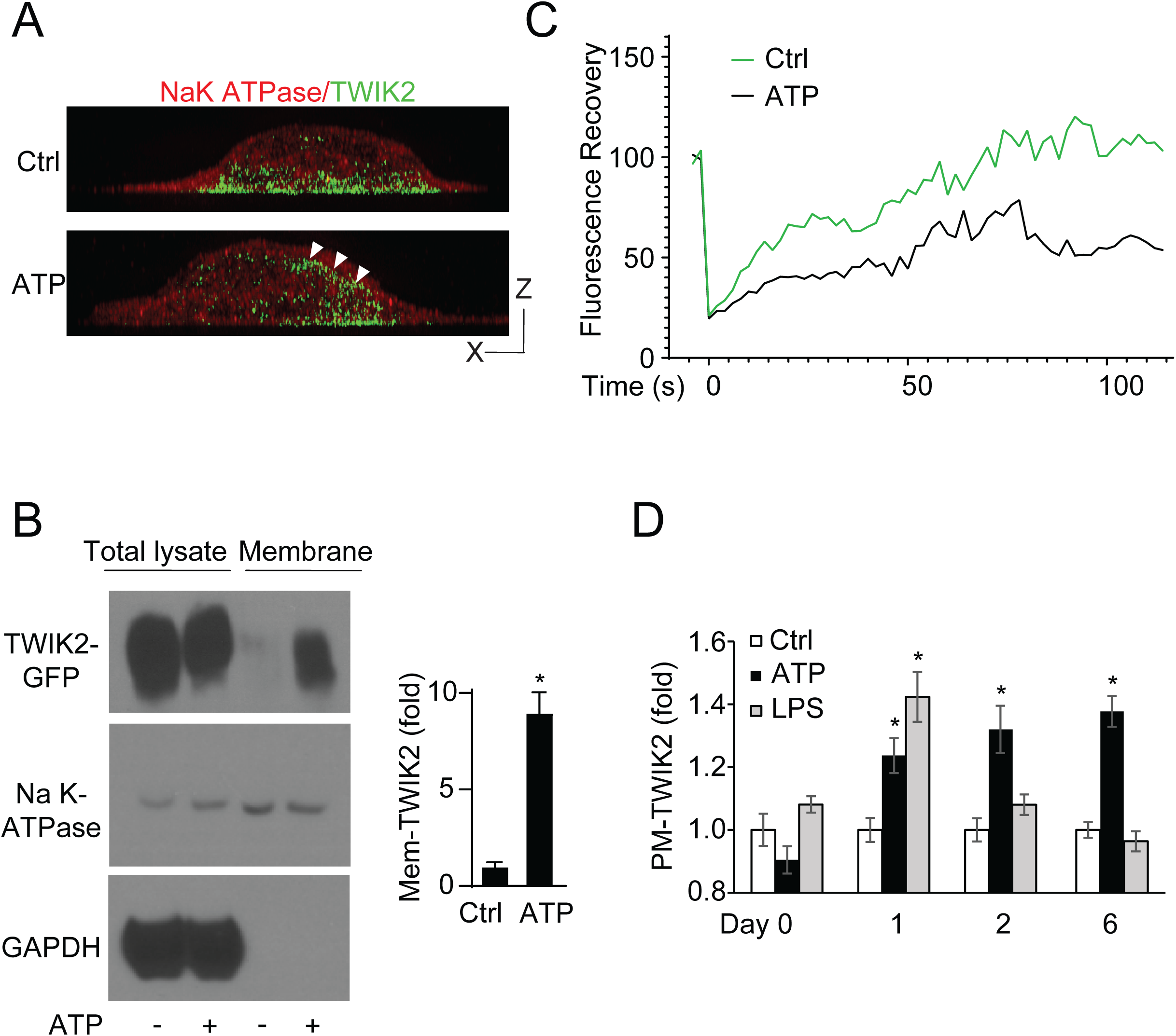
Assessment of TWIK2 PM and phagosome translocation. **A.** Images of RAW cells expressing TWIK2-GFP post-ATP challenge (1mM, PM stained in red). White arrowheads show PM translocated TWIK2. Bar, 1µm. **B.** Western blot analysis of ATP trained or untrained macrophage lysate after membrane isolation. Membrane and non-membranous fractions were compared to assess the relative abundance of TWIK2 and the Na-K ATPase with and without ATP training. **C.** Fluorescence Recovery After Photobleach of membrane-associated TWIK2-GFP in ATP-or control-treated cells. **D**. Ratiometric fluorescence intensity analysis of TWIK2-GFP signal within 1 micrometer of the plasmalemma to the corresponding TWIK2-GFP cytoplasmic intensity, highlighting enhanced and long-term membrane association after ATP treatment. Duration of PM-TWIK2 untreated or ATP or LPS treated cells is shown. *, P<0.05, compared with control. Data are representative of (A, B-left panel) or from (B-right graph, D) three independent experiments, mean and s.e.m. in B, D.

Upon phagocytosis, TWIK2 translocation to the phagosomal membrane may promote enhanced bactericidal activity through regulating phagosome acidification and ionic strength (Reeves et al., 2002). Therefore, we compared TWIK2 localization during and after phagocytosis in TWIK2-GFP expressing RAW264.7 cells trained with ATP vs. control cells. While both groups (ATP trained and untrained) efficiently endocytosed zymosan-coated beads, ATP training markedly increased TWIK2-GFP phagosomal membrane translocation as compared to untrained control cells (**Fig. 4A, B**). Thus, TWIK2 after its PM insertion is internalized in its proper topology and maintained in the membrane of particle-laden phagosomes.

**Figure 4.**
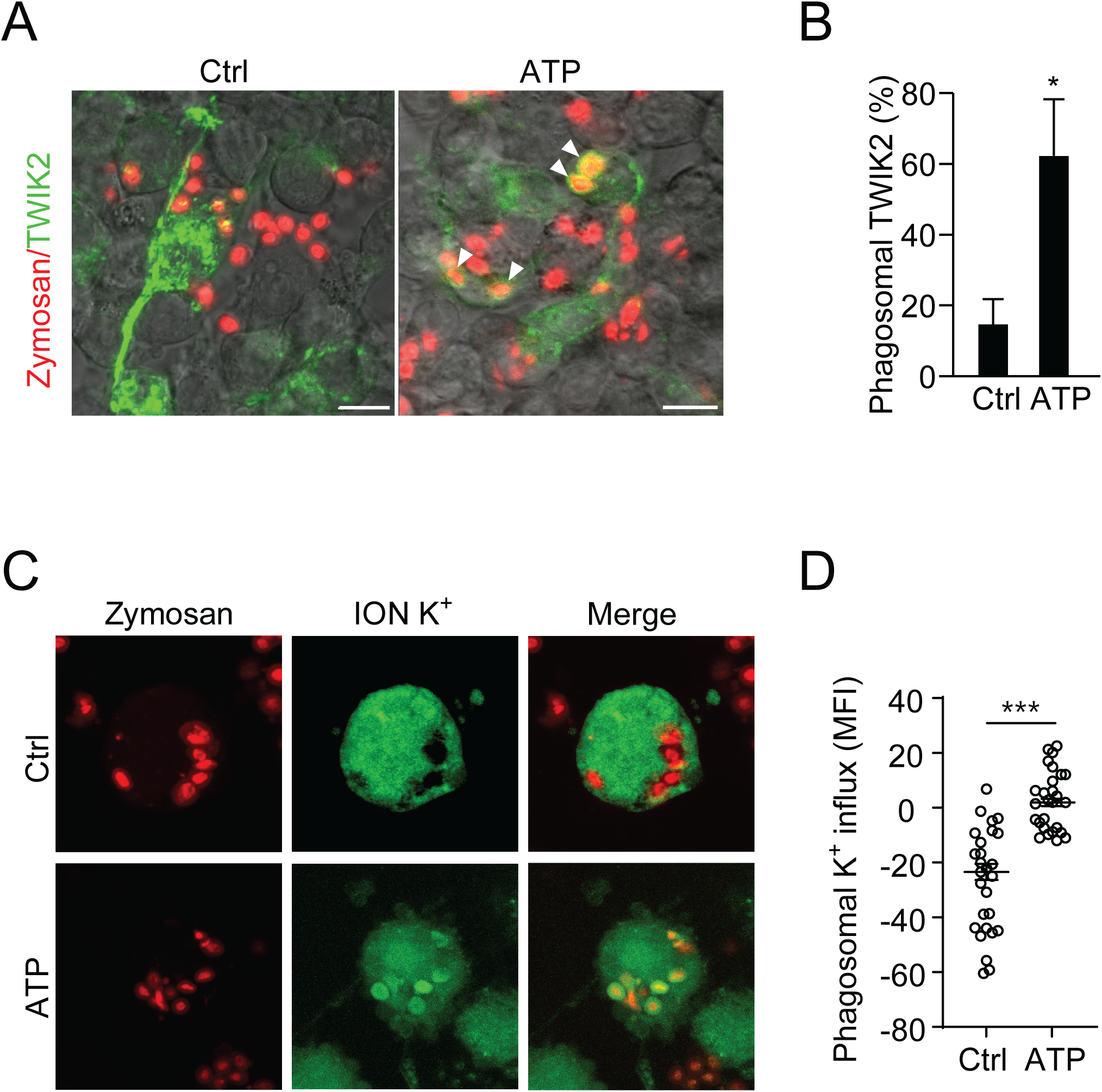
PM-TWIK2 re-internalizes into phagosomes upon phagocytosis. **A.** Fluorescence images of TWIK2-GFP expressing RAW 264.7 cells phagocytosing AF594-conjugated zymosan (red) with or without ATP treatment. **B.** Quantification of TWIK2 localization frequency in A. We observed significantly more frequent phagosome-associated TWIK2 in ATP-trained cells vs. controls. Arrowheads point to phagosomes with TWIK2-GFP. ∗, p<0.01 compared with untreated control (n = 3). **C, D**, Representative images (C) and quantification (D) of K^+^ enrichment (indicated by a specific K^+^ dye, ION K^+^, green) in phagosomes relative to cytoplasm of control and ATP-trained BMDMs. Each point (D) represents the average, net fluorescence intensity of all zymosan bead-containing phagosomes relative to the cytoplasm in one BMDM under indicated conditions (Total n= 36 Ctrl, n=45 ATP). ∗∗∗, p < 0.001. Data are representative of (A, C) or pooled from (B, D) three independent experiments, mean and s.e.m. in B, D.

Next, we compared phagosomal K^+^ changes after bone marrow-derived macrophages (BMDM) phagocytosed zymosan. We found that untrained cells exhibited little to no net phagosome K^+^ influx whereas ATP trained BMDM markedly increased net phagosome K^+^ influx relative to the cytoplasmic concentration (indicated by a specific K^+^ dye, ION K^+^, green) **(Fig. 4C, D,** dye-negative control cells were shown in **Supp. Fig. 1)**. Thus, TWIK2 translocates to phagosome membrane upon phagocytosis of trained macrophages and increases the phagosome ionic strength, which favors bacterial clearance. K^+^ influx in phagosomes via TWIK2 renders phagosomes hypertonic, which is known to activate hydrolytic proteases through the fusion of granules(Vieira et al., 2002). Indeed, ATP-induced macrophage training effects, as measured by augmented bactericidal activity, were diminished in macrophages treated with protease inhibitors **(Fig. 5A)**.

**Figure 5.**
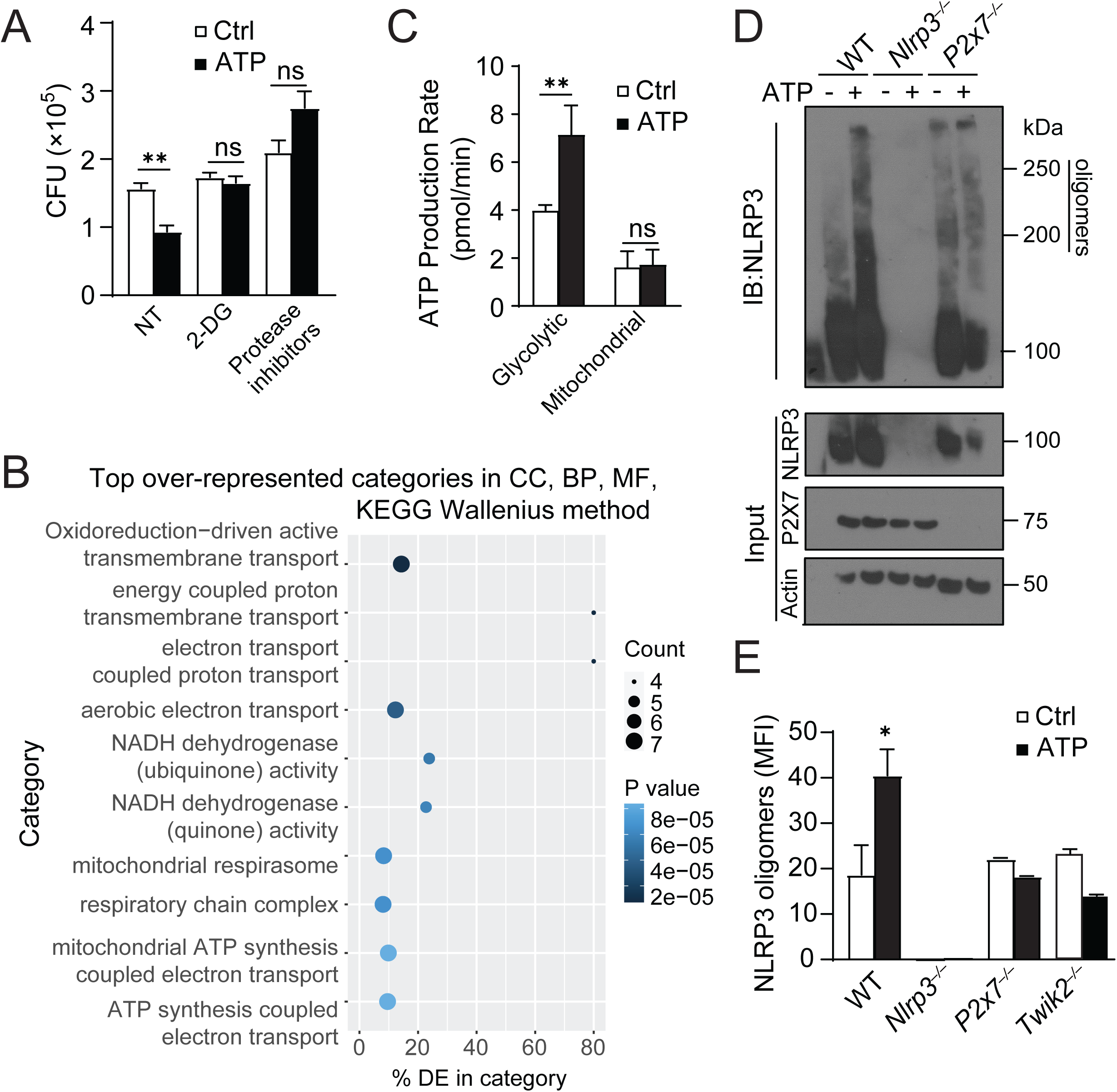
ATP training regulates transcriptional profile and metabolic reprogramming of macrophages. **A.** *PA* killing ability (as indicated by the survival of *PA*, CFU) of BMDMs trained with or without ATP in the presence of metabolic inhibitors 2DG or protease inhibitors. **B.** RNA isolated from BMDMs trained with or without prior ATP treatment was mapped to the *Mus musculus* genome and subsequently sorted into GO terms in Galaxy. Shown are the top 10 GO categories overrepresented in ATP trained BMDMs. **C.** Basal glycolytic and mitochondrial metabolism in BMDMs trained with or without ATP seeded equivalently 2 days prior to assessment as exhibited by Seahorse assay. **D, E.** Western blot (D) and quantification (E) of NLRP3 assembly in BMDMs trained with or without 1mM ATP and challenged with 1.0 MoI *PA* for 2 hours. These cells were lysed with DSP crosslinker and assessed for NLRP3 oligomerization. Data are representative of (B, D) or from (A, C, E) three independent experiments, mean and s.e.m. in A, C, E.

### ATP-TWIK2 signaling regulates transcriptional profile and metabolic reprogramming of macrophages

Since epigenetic remodeling and metabolic reprogramming are two processes that may be fundamental to the mechanisms of trained immunity (Netea and Joosten, 2018), we further examined how ATP training regulates transcriptional profile in macrophages. To identify transcriptional changes of ATP-exposed and control cells, BMDMs were trained with or without ATP, maintained in fresh media for 7 days, and evaluated by RNA-seq. GO analysis of the bulk transcriptome was performed in these trained and control cells. We discovered that the differentially expressed functional groups most markedly upregulated after ATP training are metabolic genes (**Fig. 5B**, the differentially expressed genes identified in these groups were shown in **Supp. Table 1**). To further assess the effect of the differentially regulated metabolic programs of trained and untrained macrophages, we evaluated each set via the SEAHORSE ATP production rate assay. This revealed that ATP-treated BMDMs displayed increased metabolic activity, specifically by way of glycolytic ATP production (**Fig. 5C**). Importantly, ATP-induced augmented bactericidal activity was abolished in macrophages treated with competitive metabolic inhibitor 2-deoxy-d-glucose (2-DG) (**Fig. 5A**).

Due to the dependance of bacteria killing on NLRP3 inflammasome, we assessed the role of ATP training on the NLRP3 inflammasome activation. By lysing cells in the presence of cross-linker, we assessed the extent of high molecular weight oligomerized NLRP3 and inflammasome complex formation in BMDMs with two hours of bacteria co-incubation. In untrained macrophages, exposure to bacteria resulted in the formation of 200 and 250 kDa complexes (**Fig. 5D, E**), indicating NLRP3 inflammasome assembly. This oligomerization was enhanced in ATP trained BMDMs, but was absent in P2X7 or TWIK2 knockout macrophages, in which NLRP3 oligomerization exhibited no difference between trained or nontrained P2X7, or TWIK2 null BMDMs (**Fig 5D, E**). NLRP3 is known to interact with some metabolic enzymes such as 6-phosphofructo-2-kinase/fructose-2,6-bisphosphatase 3 (PFKFB3) (Finucane et al., 2019) and phosphoglycerate kinase 1 (PGK1)(Zhu et al., 2024). Hence, ATP-mediated macrophage training may enhance metabolic activity through the NLRP3 inflammasome.

We also performed ATAC seq analysis in ATP-trained and control cells to evaluate regions of differentially accessible chromatin (**Supp. Fig. 2**). Interestingly, NLRP3 is among the genes with consistently more accessible chromatin within 2 kb of the upstream promoter region (**Supp. Fig. 2**), highlighting the importance of NLRP3 inflammasome in mediating ATP training.

### ATP training rescues pneumonia-induced immunosuppression

Bacterial pneumonia after influenza is a leading cause of severe respiratory infections worldwide (Medger and Sun, 2013). Unlike adenoviral infection (Yao et al., 2018), influenza infection abrogates macrophage-dependent bacteria clearance (Verma et al., 2020), indicating that context (type, dose and duration of infection) may be critical for macrophage phenotypes. We utilized a double-infection model to mimic this clinical scenario. Mice were first subjected to bacterial (*E. coli*) pneumonia, left to recover with or without ATP training (1 mM, i.t) for 7 days, and then infected i.t. with GFP-*PA* to cause secondary pneumonia (**Fig. 6A**). We found that the bacterial burden in lung tissue of mice doubly infected with both *E. coli* and *PA* was always greater than in mice without the primary infection, but that the extent of this bacterial load increase was markedly reduced with ATP treatment one week prior to secondary infection (**Fig. 6B**), which was not observed in other DAMP treatment such as NAD (**Fig. 6B**). This rescue effect by ATP was not observed in P2X7, NLRP3 or TWIK2 KO mice, indicate the importance of P2X7-TWIK2-NLRP3 singling in ATP mediated macrophage training (**Fig. 6C**). These results demonstrate that macrophages develop with impaired capacity to ingest or kill bacteria during pneumonia and the dysfunction can be rescued by ATP training.

**Figure 6.**
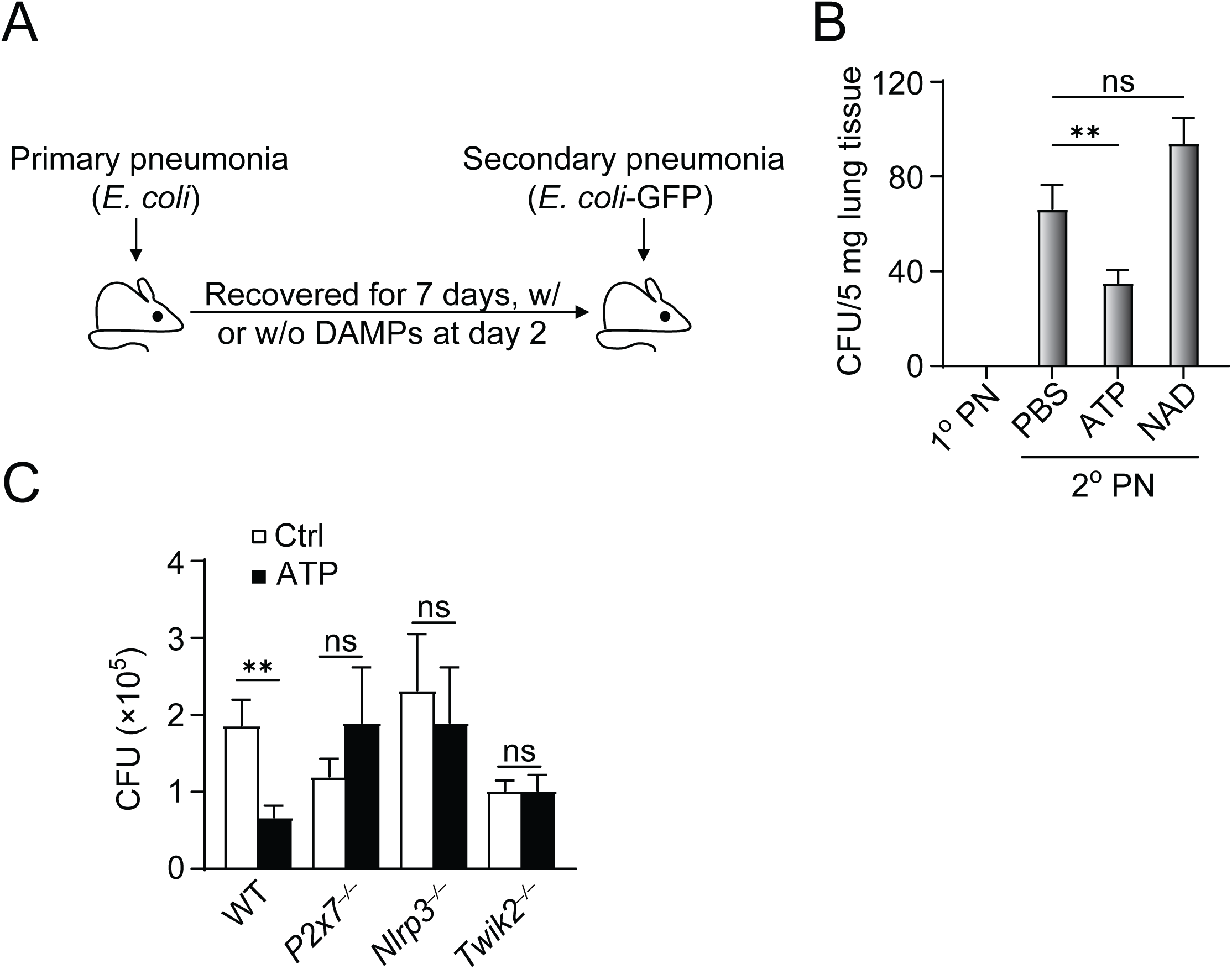
ATP training rescues lung immunosuppression caused by pneumonia. **A.** Diagram of the double-infection model of primary pneumonia (1° PN) with 1×10^6^ *E. coli* and secondary pneumonia (2° PN) with 1×10^6^ GFP-*PA*. **B.** Colony-forming units (CFU) per 5mg of lung homogenate with 1° or 2° pneumonia. ∗, p < 0.05. **C.** Relative bacterial killing ability in lung homogenate of WT or designated KO mice trained with or without ATP in the double exposure model, n = 5. Data are from (B, C) three independent experiments, mean and s.e.m. in B, C.

## Discussion

Lungs are continuously exposed to environmental pathogens and require a rapid immune response to ensure host survival. Lung resident macrophages such as AMφ are key cells of this innate defense, and induction of trained innate immunity in these cells enhances their capability. Leveraging the trained immunity of macrophages would be beneficial to promote host defense function while preventing auto-inflammatory injury, the key feature of ALI/ARDS. We discovered that ATP could train macrophages through the ATP-P2X7-TWIK2 pathway described above. Our previous data suggest that during homeostasis, TWIK2 resides in the endosomal compartment (Huang et al., 2023). In response to increased extracellular ATP, endosomal TWIK2 is translocated to the PM and remains localized in the phagosome membrane upon phagocytosis of bacteria. This raises the intriguing concept of augmenting the activity of TWIK2 that will train macrophages in multiple ways. We posit that a noteworthy element of this function is the tunability of TWIK2 association with PM and phagosomes and hence the ability of TWIK2 localization to control innate immunity for a relatively long duration (the definition of training). We hypothesize that tunability of macrophages is a central function of these cells and it requires upregulation and/or duration of ATP-mediated TWIK2 PM translocation. Therefore, controlling the magnitude and duration of TWIK2-mediated NLRP3 inflammasome activation provides potential therapeutic targets for the treatment of ALI.

Our data showed other DAMPs such as histone or NAD did not elicit trained immunity of macrophages indicating the importance of ATP signaling. Different pathogens or their products exploit distinct mechanisms to train macrophages. For example, adenoviral infection induced macrophage trained immunity requires IFN-γ produced by T-cells (Yao et al., 2018), while LPS-induced AMφ training depends on type 1 interferon, IFN-β (Zahalka et al., 2022). LPS trained AMφ also exhibited upregulated expression of the efferocytosis receptor MERTK and greater capacity for efferocytosis of cellular debris (Chakraborty et al., 2023). Thus, generalizations may not be possible. In addition, since viral, LPS or bacterial challenge can damage other nearby tissue and therefore generate ATP, it will be interesting to test whether ATP-TWIK2 contributes to different means of training.

There is the possibility that training and suppression are both induced upon different pathogen challenges (Verma et al., 2020; Yao et al., 2018). Both the nature of the pathogen and the pathogen dose play critical roles in determining these opposing effects (Bauer et al., 2018). For example, low-dose LPS promotes hyperactive responses while high-dose LPS prolongs inhibition in macrophages (Bauer et al., 2018; Fu et al., 2012; Zahalka et al., 2022). In addition, trained innate immunity is different from a primed immune response. In contrast to priming, trained immunity returns to the basal level following removal of the first stimulus (Divangahi et al., 2021). However, in response to secondary challenges, both gene transcription and cell functions are enhanced at much higher levels than those in the primary challenge (Divangahi et al., 2021). In our training model, the functional program of macrophages returns to basal level after first stimulation. The training model is fundamentally different from models of priming in which the stimulus is either maintained for a long period of time to induce priming or secondary stimulation is performed very quickly after the initial priming.

This study has several limitations, one of which concerns the specificity and causal interpretation of the observed phenotypes. Although mechanistic experiments were performed in RAW264.7 cells and primary BMDMs, these findings were extrapolated to the *in vivo* setting primarily through correlations with bacterial clearance outcomes. Consequently, we cannot exclude the possibility that other tissue-resident immune populations also respond to eATP in ways that contribute to the enhanced bacterial control observed *in vivo*. In addition, we have not directly assessed potential deleterious consequences of eATP-mediated immune activation. Nonetheless, the improved survival in ATP-treated mice with *PA* pneumonia, together with the comparable magnitude of bacterial clearance in ATP-treated macrophages *in vitro* and *in vivo*, suggests that these limitations do not undermine the central conclusion that eATP augments macrophage-associated antibacterial responses. The precise contribution of macrophages relative to other immune cell types, as well as any associated risks of eATP-enhanced immunity, will require further investigation.

The discovery of trained immunity has profound implications for both immunity to infectious diseases and the development of novel therapeutic strategies. Despite the benefits, there are concerns about the potential harmful effects of a chronically trained, or partially activated, immune system (Netea et al., 2016). Thus, understanding the mechanisms that govern the induction, persistence, and resolution of trained immunity is essential for developing strategies to harness its benefits while minimizing its risks. Our findings reveal that regulatable association of TWIK2 with the PM and phagosomes triggered by ATP mediates trained immunity in macrophages. The tunability of macrophages is a central function of these cells and it requires upregulation and/or increased duration of ATP-mediated TWIK2 PM translocation, which will lead to understanding new immunotherapeutic approaches.

## Methods

### Cell Culture and Mice

The RAW 264.7 mouse macrophage cell line was obtained from ATCC (TIB-71). RAW 264.7 cells were cultured in Dulbeccoʹs Modified Eagle Medium (DMEM) supplemented with 10% fetal bovine serum (FBS) and 1% penicillin-streptomycin and were maintained at 37°C in a humidified atmosphere with 5% CO₂.

For in vivo tracking of TWIK2, RAW264.7 cells were transfected with TWIK2-EGFP plasmid (pLV[Exp]-Puro-CMV-mKcnk6 [NM_001033525.3] /3xGS/EGFP), designed by and purchased from VectorBuilder (VB200618-1166ypg) and cultured in puromycin selection media. Transfected cells were further selected by GFP expression. Bone marrow-derived macrophages (BMDMs) were isolated from the femurs and tibias of C57B/L6J or C57B/L6J -derived mice (8-16 weeks old, males and females harvested approximately equally) and differentiated by culturing in RPMI 1640 with 10% FBS and 1% penicillin-streptomycin supplemented with 0.22 μm-filtered 10% L929-conditioned media for 5 days. BMDMs were cultured in RPMI 1640 with 10% FBS and 1% penicillin-streptomycin. These cells were also maintained at 37°C in a humidified atmosphere with 5% CO₂ for the duration of the experiment.

All mice were handled in accordance with NIH guidelines for the care and use of laboratory animals and UIC animal care and use committee (ACC)’s regulations. All procedures were approved by the UIC IACUC. Mice of 8–16 weeks of age, body weight ranging from 18–25 g, were used for all experiments. P2rx7^−/−^, Nlrp3^−/−^ mice were purchased from Jackson Laboratory. Myeloid knockdown of TWIK2 was generated by maintaining heterozygous LysM^cre^ in a TWIK2^f/f^ background. Deletion of specific genes was confirmed by PCR and western blot.

### ATP-mediated training

For ATP training in cell culture, ATP salt (Thermofisher L14522.06) was dissolved at 1.0M in cell culture-grade water. This stock was diluted to a final concentration of 1mM in complete RPMI media, warmed to 37°C and subsequently applied to cells. Media was changed and cells were washed in PBS to further dilute remaining ATP after 24 hours, and cells were maintained for an additional 6 days until challenge or harvest. Bacterial cultures for challenge (*E. coli* strain DH5a and/or GFP-transformed *P. aeruginosa* strain PA-01) were grown in Luria Broth or Luria Broth plus 0.1 mg/mL Ampicillin, respectively, for 12-16 hours. Culture density was determined by empirically determined OD600 and bacteria (or AF594-tagged zymosan beads (Thermofisher Z23374)) were diluted to 2.5×10^7^ CFU (colony forming units) or beads per mL and co-cultured with cells in culture conditions described above. Primary challenge (*E. coli*) was co-cultured in complete RPMI 1640 media for 24 hours before washing with PBS and replenishing sterile complete media. *P. aeruginosa* and Zymosan bead endpoint challenge was co-cultured in sterile PBS for 2 hours (phagocytosis, bacterial clearance, immune cell recruitment, NLRP3 assessment, RNA-seq), or 24 hours (ELISA) before final assessment. TWIK2-GFP localization, and ATAC-seq received no endpoint challenge to avoid confounding changes to gene expression.

### Tissue preparations for single-cell suspension and flow cytometry

Whole lungs were harvested, minced, and enzymatically digested in PBS containing collagenase type IV (1 mg/ml), hyaluronidase (0.15 mg/ml), and DNase I (0.05 mg/ml) at 37°C for 30 min in a shaking incubator. All enzymes were purchased from Sigma-Aldrich. Digested tissues were passed through a 70-µm cell strainer to obtain single-cell suspensions. After washing with PBS, red blood cells were lysed with BD Pharm Lyse Lysing Buffer (BD Bioscience) for 10 min at 4°C. The remaining cells were washed and counted using an automated cell counter.

For flow cytometric analysis, single-cell suspensions were stained with fluorochrome-conjugated antibodies against CD11c (BV605), CD64 (APC), CD11b (APC-Cy7), CD45 (PerCP-Cy5.5), Siglec-F (PE), Ly6C (PE-Cy7), and Ly6G (BV510) for 20-30 min on ice. All antibodies were purchased from BioLegend (San Diego, CA, USA). Following staining, cells were washed with PBS and acquired on an LSR Fortessa flow cytometer (BD Biosciences). Flow cytometry data were analyzed using FlowJo software (v10.10.1).

### Cytokine quantification

Bronchoalveolar lavage fluid (BALF) was collected, followed by centrifugation at 400 × g for 10 min at 4 °C to remove cells. Supernatants were aliquoted and stored at −80 °C until analysis. Concentrations of IL-1β, IL-18, and IL-6 were quantified using Quantikine® ELISA kits (R&D Systems) according to the manufacturer’s instructions. Briefly, non-tissue culture treated 96 well plates were incubated with capture antibody overnight. BALF samples were thawed on ice, and subsequently incubated in coated 96-well plates. After washing, wells were incubated with horseradish peroxidase–conjugated detection antibodies, followed by substrate development with stabilized TMB. Absorbance was measured at 450 nm with wavelength correction at 540 nm using a microplate reader. Cytokine concentrations were calculated from plate-controlled standard curves and expressed as pg/mL.

### Survival Study

C57B/L6J mice were anesthetized and intranasally treated with 40 μL of cell culture-grade sterile water containing the indicated amount of dissolved ATP salt. After 6 days, mice were again anesthetized to be intranasally inoculated with GFP-expressing *Pseudomonas aeruginosa* (strain PA-GFP-01) at a concentration of 5×10^6^ CFU in a total of 40 μL cell culture-grade sterile water while in a BSL2 facility. 7 animals per group were used for the survival studies.

### Bacterial Killing Assay

An in vitro bacterial killing assay was conducted using ampicillin-resistant *P. aeruginosa* (strain PA-01 GFP). Macrophages were co-cultured with bacteria at a multiplicity of infection (MOI) of 1.0. After 2 hours of incubation, samples of the supernatant were assessed to evaluate remaining bacterial load.

Macrophages were washed with sterile PBS, briefly incubated with gentamicin, and lysed with cell culture grade water. Serial dilutions of both samples were plated on LB agar with 0.1mg/mL Ampicillin to quantify bacterial load (supernatant) and killing (lysate). Bacterial colonies were counted after overnight incubation at 37°C. To assess in vivo bacterial clearance/bacterial load, mice were anesthetized and treated intranasally as above with ATP and, later, *P. aeruginosa.* Lungs were harvested 2 hours after inoculation, weighed, and homogenized. Homogenate was then plated on selective media to assess CFU per lung mass.

### Microscopy

Structured illumination microscopy (SIM) was performed using an DeltaVision OMX 3D-SIM microscope (GE Healthcare) with 3D-SIM reconstruction software, using a 60×, 1.42 NA Oil PSF Objective (Olympus PlanApo N). Wild type BMDMs were stained with ANTIBODIES and imaged. Membrane GFP intensity was measured using a Leica LSM710 BiG confocal microscope with a Pecon XL TIRF S incubation system for long term imaging of cultured cells. TWIK2-GFP expressing RAW cells were imaged with a 63× Plan-Apochromat oil objective, 1.4 NA objective once daily, and the TWIK2-GFP fluorescence intensity within 1 micron of the plasma membrane was ratiometrically compared to the average TWIK2-GFP intensity of the measured cell with ImageJ software version 1.54g.

### Membrane Isolation

Plasma membranes were isolated using the Subcellular Protein Fractionation Kit for Cultured Cells (Thermo Fisher Scientific 78840) according to the manufacturer’s instructions, using the maximum recommended 10×10^6^ trained or untrained BMDMs per sample. Fractionation was verified as described by Thermo Scientific for the Sorvall MTX Micro-Ultracentrifuge and S55-A2 Rotor. Briefly, post-nuclear lysate of ATP-trained and untrained cells was layered on a 0-30% isotonic percoll stepwise gradient and separated via ultracentrifugation at 84000×g for 30 minutes at 4°C, the membrane-containing lysate was collected from the interface, diluted in ice-cold PBS, and further ultracentrifuged at 105,000×g for 90 minutes at 4°C to clear percoll and obtain purified membranes.

### In vitro Chemical challenge and assessment

To evaluate the role of metabolism on ATP effect, metabolic inhibitors 2-Deoxy-D-glucose, NG52, and PFK158 (All from SelleckChem) were applied to cells at 1mM, 10 μM, and 15 μM respectively in incomplete media (not supplemented with FBS or antibiotics) for 1 h (NG52, PFK158) or 3 h (2DG) pretreatment. *P. aeruginosa* was then applied to culture to assess bacterial clearance as described above. To assess TWIK2 dynamics during phagocytosis, 0.1 mg of AF594-conjugated zymosan particles (Thermofisher Z23374) were added to vehicle- or ATP-treated TWIK2-GFP transfected RAW264.7 cells (approximately 2 particles per cell). Positively GFP expressing cells were imaged and assessed for 1) engulfment of particles and 2) TWIK2 localization. Frequency of engulfed particles that colocalize with TWIK2-GFP was reported.

ION Potassium Green (working concentration 5 µM in DMSO; Abcam AB142806) or equivalent volume DMSO vehicle was applied to cells concurrently with zymosan beads and assessed by confocal microscopy as described above. To assess net phagosomal K^+^ influx, the baseline, AF594^-^, cellular area green fluorescence was subtracted from that of the AF594^+^ region, with negative values indicating higher green MFI and thus low phagosomal K^+^ influx.

### Fluorescence Recovery After Photobleaching (FRAP)

FRAP was performed to assess the dynamics of TWIK2-GFP on phagosomal membranes. Cells were bleached using a 488 nm laser at maximum laser power, and recovery was monitored for 2 minutes with images captured every 2 seconds, using an 63x Plan-Apochromat oil objective, 1.4 NA. Analysis was conducted using ImageJ software version 1.54g, and percent recovery was determined by ratiometrically comparing each measurement to the average of two distinct pre-bleach measurements.

### Seahorse ATP Production Assay

ATP production was quantified using a Seahorse XFe 96 Analyzer (Agilent Technologies). BMDMs were differentiated and trained in 60 mm dishes, detached via Accutase 48 hours prior to assessment, live cells counted by a Typan-blue adjusted Countess II automated cell counter, and seeded in a Seahorse XFe96 cell culture microplates at 5×10^4^ cells per well. Rotenone/Antimycin and Oligomycin A were loaded per manufacturer’s instructions, and basal and maximal respiration rates were measured, according to the manufacturer’s protocol for the XF Real Time ATP Rate Assay Kit.

### NLRP3 Oligomerization Assay

Cells were washed with cold, sterile PBS and lysed in RIPA buffer (western blotting) or NP40 buffer with 1 mM DSP crosslinker (NLRP3 oligomerization). For NLRP3 oligomerization experiments, lysates were centrifuged at 10,000 r.p.m. for 20 min to pellet the insoluble fraction and insoluble lysates were washed and diluted into NP-40 lysis buffer. Approximately equivalent amount of protein was loaded and separated on 8–12% polyacrylamide gels and transferred onto activated PVDF membranes. Membranes were blocked in 5% skim milk then incubated with primary antibodies and secondary horseradish peroxidase (HRP)-conjugated antibodies. Antibodies used were anti-Nlrp3 (Adipogen, AG-20B-0014-C100), and anti-TWIK2 (LS Bio LS-C110195-100). Actin, GapDH, or ATP1A1 were used to verify loading accuracy as indicated. Blots were developed on Xray film and scanned at 300 dpi. Intensities/blots were otherwise unmodified. Band intensities were quantified using ImageJ by selecting rectangular regions of interest around each band and subtracting background signal from adjacent areas.

### RNA-seq analysis

Analysis was adapted from Reference-based RNA-Seq data analysis (Galaxy Training Materials, https://training.galaxyproject.org/training-material/topics/transcriptomics/tutorials/ref-based/tutorial.html). In brief, RNA was isolated from ATP-treated and vehicle-treated populations of BMDMs via a Qiagen RNeasy mini kit. Concentration was normalized and raw paired-end RNA-seq reads were obtained in FASTQ format via an Illumina NovaSeqX [Uchicago genomics facility]. Sequences were uploaded to a Galaxy server and processed therein. Quality control was performed using FastQC, and adapter trimming was conducted with CutAdapt using default parameters. Trimmed reads were aligned to the reference genome using RNA STAR. Genome and annotation files were retrieved from Ensembl and indexed prior to alignment. Aligned reads were quantified using featureCounts to generate gene-level count matrices. Differential expression analysis was performed using DESeq2, with significance thresholds set at adjusted p-value < 0.05. Gene Ontology enrichment analysis was conducted using goSeq to assess biological relevance. Raw RNA sequencing data submitted to the NIH sequence read archive - PRJNA1356819.

### ATAC-seq analysis

Raw reads were mapped to the mouse reference genome (mm39) using BWA MEM (Li, 2013) with default parameters. Apparent PCR duplicates were removed using Picard MarkDuplicates. Read alignments were adjusted to account for Tn5transposon binding: +4bp for + strand alignments, -5bp for - strand alignments following the guidelines for the ATAC-seq protocol as published (Buenrostro JD, 2015). The open chromatin enrichment track was generated by creating a bedGraph from the alignments BED file using bedtools genomcov (Quinlan and Hall, 2010), then converted to bigWig using UCSC tool bedGraphToBigWig (Kent et al., 2010); tracks were normalized by the sum of total aligned read lengths over 1 billion. Open chromatin peaks were called using Macs2 (Zhang et al., 2008) with --nomodel set and no background provided; peaks with a score >5 were retained.

For differential open chromatin analysis, differential analysis of quantitated peaks as compared with treatment was performed using the software package edgeR on raw peak counts (McCarthy et al., 2012). Data were normalized as counts per million and an additional normalization factor was computed using the TMM algorithm. To account for different processing batches, a batch-correction factor was added to the model for the analysis. Statistical tests were performed using the “exactTest” function in edgeR. Adjusted p values (q values) were calculated using the Benjamini-Hochberg false discovery rate (FDR) correction (Benjamini and Hochberg, 1995). Significant peaks were determined based on an FDR threshold of 5% (0.05).

For pathway enrichment analysis, gene sets for pathway enrichment analysis were obtained from promoter-associated open chromatin, with the top 400 up- and top 400 down-regulated peaks by p value included in the analysis. A core analysis targeting upstream regulators and pathway enrichment based on this gene list was performed using Ingenuity Pathway Analysis (IPA; QIAGEN Inc, https://digitalinsights.qiagen.com/IPA).

For motif enrichment analysis, instances of known transcription factor motifs were searched for in called peak sequences with comparison to the JASPAR database (Khan et al., 2018) using FIMO (Bailey et al., 2009). Motif enrichment statistics were computed for the top 1000 up- and top 1000 down-regulated peaks using Fisher’s Exact Test. This was repeated for all motifs, correcting for multiple testing using the FDR correction of Benjamini and Hochberg (Benjamini and Hochberg, 1995).

The ATAC seq has been submitted to the NIH SRA at SUB15765984.

### Statistical analysis

All experiments were performed at least three times. Graphs were generated using GraphPad Prism. All data were expressed as mean ± standard error of the mean. Statistical comparisons were made using ordinary one-way ANOVA multiple comparisons with Prism 6 (GraphPad). Number of samples or mice per group, replication in independent experiments and statistical tests are mentioned above or in the figure legends. Significance between groups was labeled with asterisks indicating a statistically significant difference.

## Acknowledgements

We would like to thank Jason M. Wood and Mark Maienschein-Cline at Research Informatics Core, Research Resources Center, University of Illinois Chicago for help with the ATAC seq analysis. Bioinformatics analysis in the project described was provided by the UIC Research Informatics Core, supported in part by NCATS through Grant UM1TR005438, and the UIC Cancer Center Cancer Bioinformatics Shared Resource. This study is supported in part by National Institute of Health grants P01HL151327 and R01DK126753 (K.K); National Natural Science Foundation of China 82130001 and Shanghai Municipal Science and Technology Major Project ZD2021CY001 (Y.S). J.T and J.C were supported by NIH T32 HL007829.

## Figure Legends

**Supplemental Table 1.** Expression levels of all differentially expressed genes and gene membership within the top 10 most significantly enriched GO categories This table reports the normalized expression values for all genes identified in the analysis. Genes making up the ten most significantly enriched Gene Ontology (GO) categories are also listed.

---

**Supplementary Figure 1.**
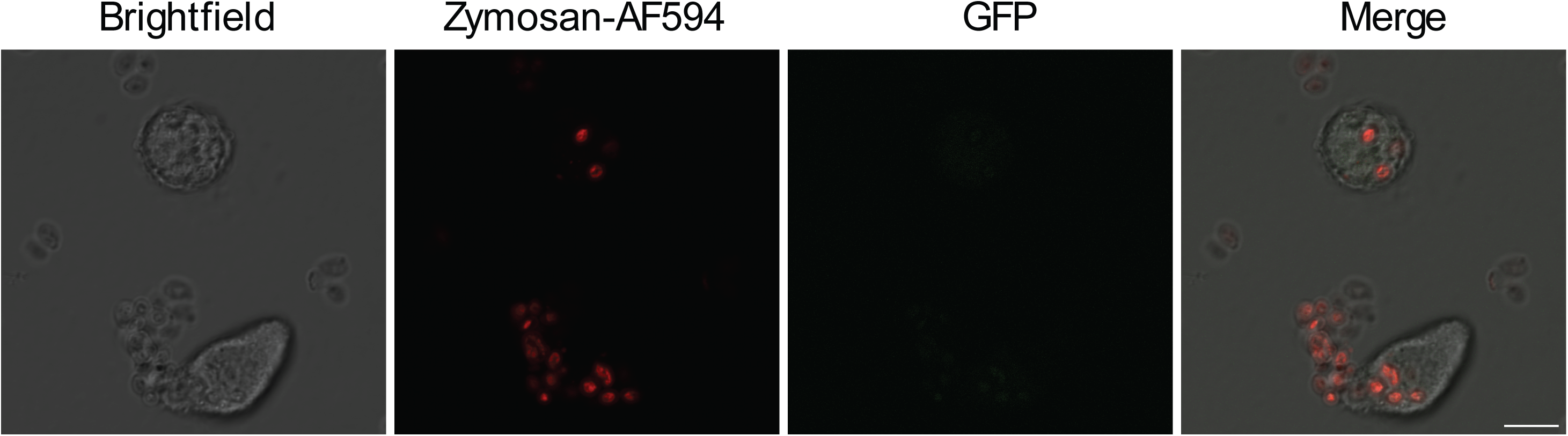
Negative controls for K^+^ enrichment as shown in **Figure 4C**. Cells were incubated with fluorescently labeled zymosan 594 beads under identical conditions in **Figure 4C**, but without addition of Ion K^+^ Green dye. Images were captured using identical laser power and selected focal plane with most up taken AF594-Zymosan beads in frame. Scale bar, 10 µm.

**Supplementary Figure 2.**
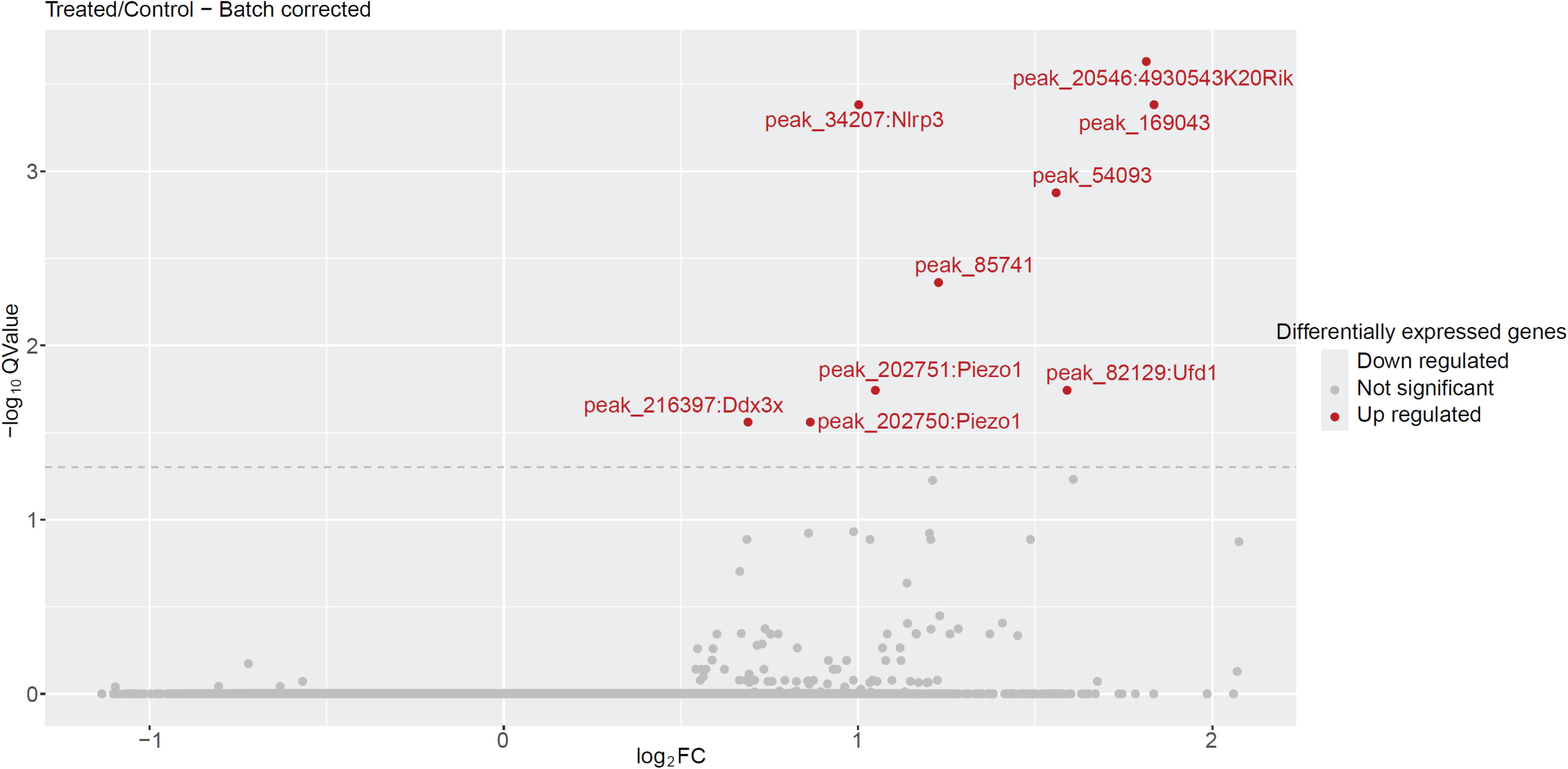
Volcano plot displaying log_2_ fold change of differentially accessible chromatin in ATP treated BMDMs relative to that of vehicle-treated BMDMs 6 days after treatment. The x-axis shows log_2_ (fold change), and the y-axis denotes - log_10_(Q-value). Peaks with Q-value (Benjamini-Hochberg FDR) are highlighted in red and labeled with the gene containing the nearest likely promoter region. No significant down-regulated peaks were noted in this comparison. The horizontal dashed line indicates the Q-value threshold of 0.05 (-1.3 on the -log_10_ scale). The ATAC-seq data has been submitted to the NIH SRA at SUB15765984.

## Notes

### Competing Interest Statement

The authors have declared no competing interest.

### Summary of Updates

This version of the manuscript has been revised to address reviewer's comments.

